# Advanced whole genome sequencing and analysis of fetal genomes from amniotic fluid

**DOI:** 10.1101/173807

**Authors:** Qing Mao, Robert Chin, Weiwei Xie, Yuqing Deng, Huixin Xu, Rebecca Yu Zhang, Quan Shi, Erin E. Peters, Natali Gulbahce, Zhenyu Li, Fang Chen, Radoje Drmanac, Brock A. Peters

## Abstract

Amniocentesis is typically performed to identify large chromosomal abnormalities within the fetus. Here we demonstrate that it is feasible to generate an accurate whole genome sequence (WGS) of a fetus from an amniotic sample. DNA from cells and the amniotic fluid were isolated and sequenced from 31 amniocenteses. Concordance of variant calls between the two DNA sources and with parental libraries was high. Two fetal genomes were found to harbor potentially detrimental variants in *CHD8* and *LRP1*, variations in these genes have been associated with Autism Spectrum Disorder (ASD) and Keratosis pilaris atrophicans, respectively. We also discovered drug sensitivities and carrier information of fetuses for a variety of diseases. In this study, we demonstrate for the first time the sequencing of the whole genome of fetuses from amniotic fluid and show that much more information than large chromosomal abnormalities can be gained from an amniocentesis.

## Introduction

Amniocentesis is a common procedure performed on over 200,000 women a year in the United States alone. It is currently performed on women who are considered to be at a higher risk for pregnancy complications because of their advanced age or to further investigate an abnormal blood or ultrasound test result (*e.g.*, suspected Down syndrome as a result of an extra chromosome 21 detected by noninvasive prenatal testing (NIPT)). The procedure involves the insertion of a needle through the wall of the uterus and the amniotic sac to collect approximately 20 ml of amniotic fluid. Cells from the fluid are collected through centrifugation, cultured, and after approximately two weeks analyzed by fluorescent in situ hybridization (FISH) or a microarray to detect abnormal chromosomal copy number changes or large chromosomal structural rearrangements. In some cases a small number of specific genes are examined for single to multi-base changes. These tests have become the gold standard for detecting Down syndrome and several other serious birth defects because they have a low false positive rate, however they are unable to detect the majority of birth defects. Currently, there are over 5,000 genes associated with a genetic disease for which a gene test is offered according to GeneTests (www.genetests.org). In addition, recent studies have defined large sets of genes in which coding variants within these genes are associated with Autism (Michaelson et al. 2012; O’Roak et al. 2012; Sanders et al. 2012; De Rubeis et al. 2014; Iossifov et al. 2014), severe intellectual disability (Gilissen et al. 2014), and other congenital disabilities (de Ligt et al. 2012; Veltman and Brunner 2012; Epi et al. 2013; Yang et al. 2013; Al Turki et al. 2014; Fromer et al. 2014; McCarthy et al. 2014; Purcell et al. 2014; Deciphering Developmental Disorders 2015) and a large population study (Lek et al. 2016) discovered thousands of genes that are intolerant to coding variants, adding to the set of genes that could be disease causing. Taken together these data provide strong evidence that there are probably thousands of genes for which coding changes or complete loss of function of the gene product are incompatible with life or could result in a serious disease phenotype. This suggests testing for large chromosomal changes or examining just a few genes is insufficient for detecting most disease causing genetic defects.

In this study, we developed a modified workflow to enable the WGS analysis of cell-free DNA (cfDNA) from amniotic fluid. For each amniotic sample, we isolated DNA from both the amniotic fluid and the cell pellet. We also collected DNA from the blood of each parent. WGS libraries were made and sequenced for each DNA sample enabling a rich set of genomic data for reproducibility analyses and clinical variant annotations to be performed.

## Results

### Determination of quality of fetal genome data

28 cfDNA and all 31 cell pellet DNA samples yielded high quality data. Coverage across each sample was excellent with a confident call for both alleles made for ~97% of the genome and ~98% of the exome from both DNA sources (Fig. 1a and b and Table S1). For about half of the cell pellet DNA samples an LFR (Peters et al. 2012) library could successfully be made. These libraries also showed good coverage with an average of ~96% of the genome and exome called confidently. Approximately 4 million variants per library were called, similar to previous studies on Asian genomes (Genomes Project et al. 2015) (Fig. 1b). For most amniocentesis samples a standard library was made from both the cfDNA and the cell pellet enabling comparisons of variant calls between each library (Table S2). In general, over 96% of calls were shared between pairwise comparisons at all locations for which both libraries were covered with sufficient reads (Fig. 2a). Additionally, since both parents were sequenced the fetal genome from each library could be compared with parental calls at each loci as further confirmation that the correct variants are being called at all positions. This showed that ~99% of calls were consistent with variant calls made in the parents (Fig. 2b). Taken together these results suggest that high quality fetal genomes can be generated from either cfDNA in the amniotic fluid or high molecular weight DNA isolated from the cell pellet.

**Figure 1.**
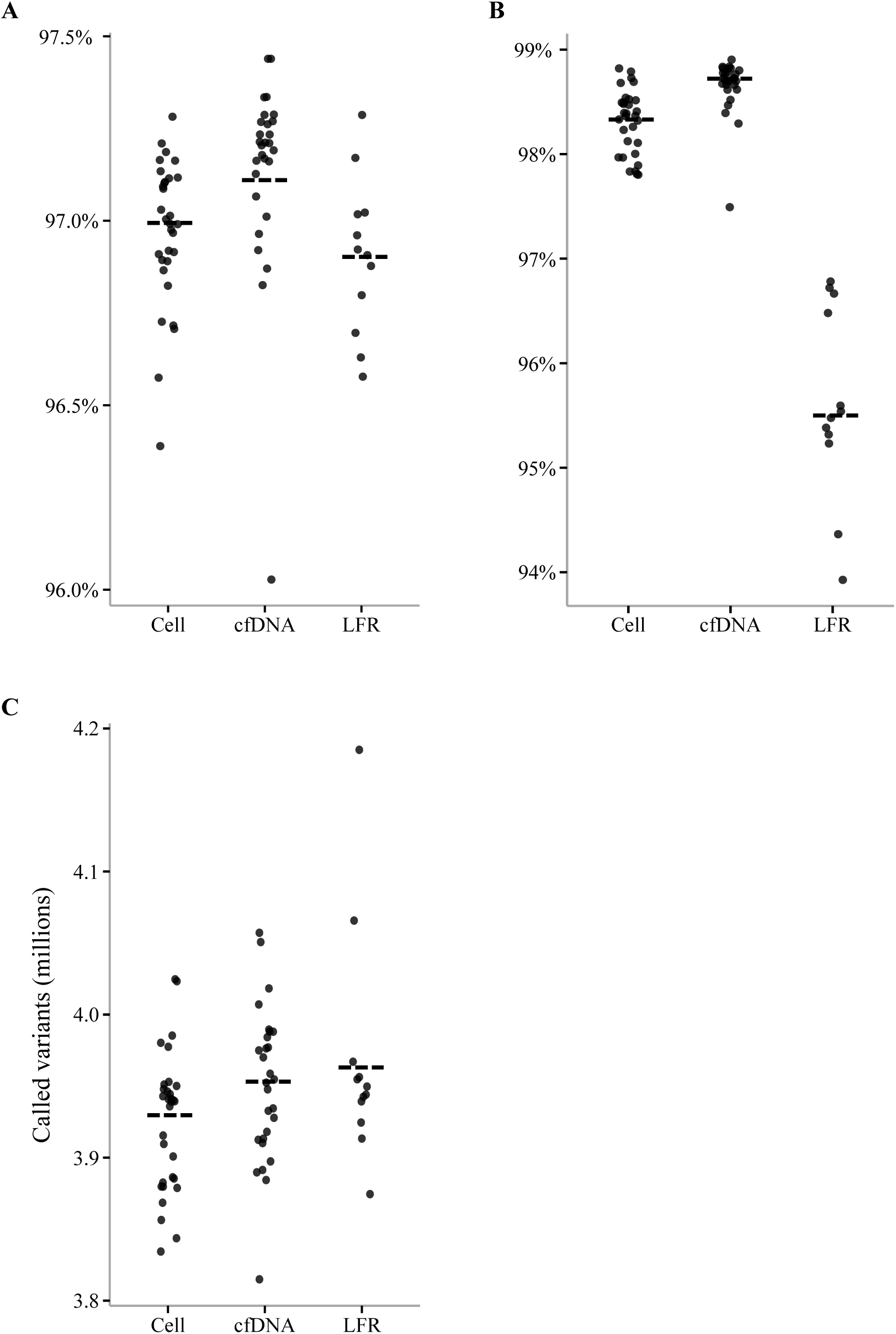
Genome and variant call performance. The percent of the genome (**A**) and the exome (**B**) called for both alleles was calculated for each library. (**C**) The total number of SNPs for each library is plotted. The red dashed line in all figures is the average rates seen in Complete Genomics’ whole genome sequencing data.

**Figure 2.**
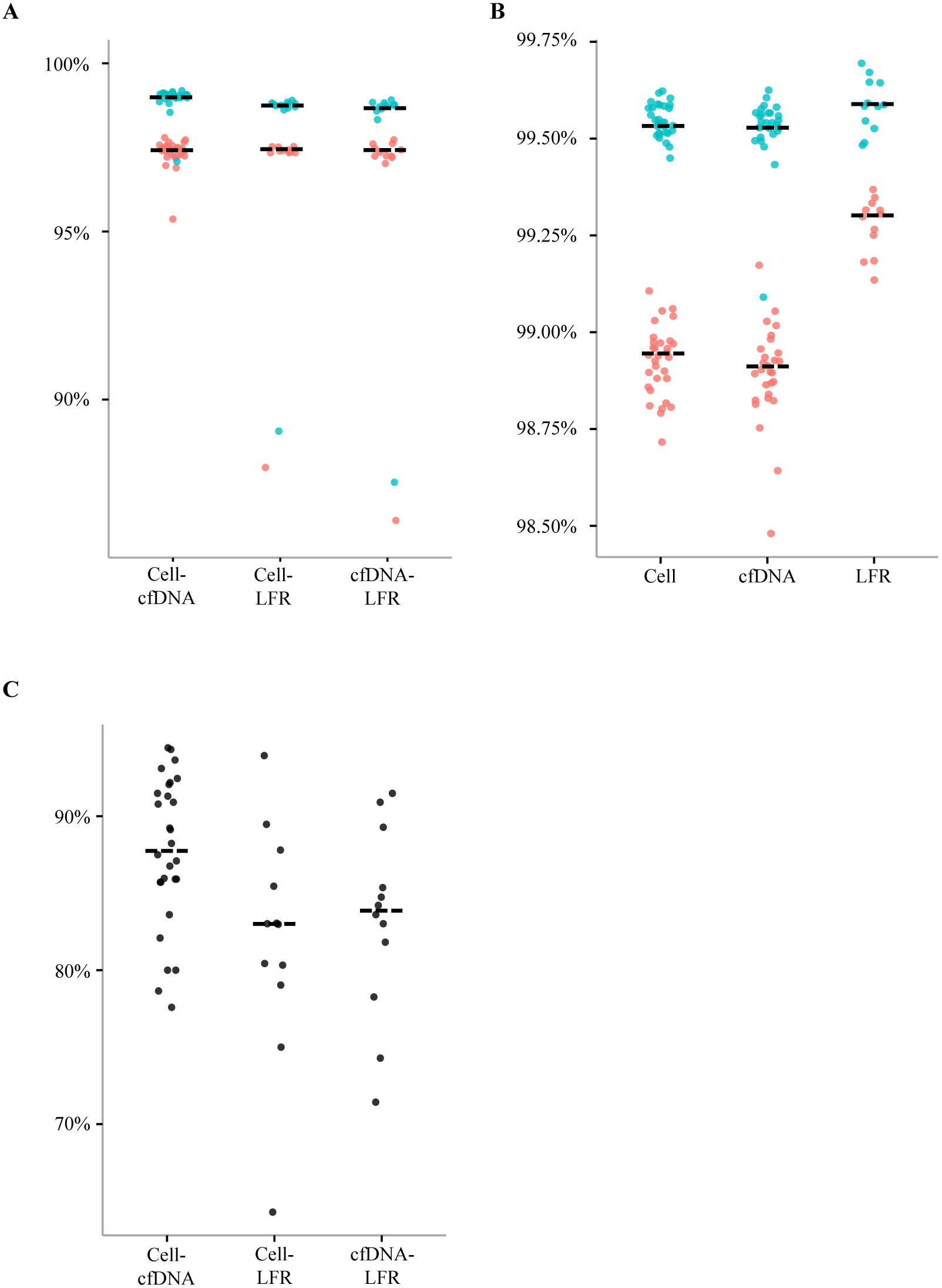
Small variant library concordance. Percent overlap between replicate samples (**A**) and the parents and fetus (**B**) are plotted. Each dot represents a pairwise comparison. Blue dots are SNPs only and red represent all variants. The overlap of DNMs between replicate samples (**C**) is also plotted.

### DNM analysis

Previous studies (Gilissen et al. 2014; Peters et al. 2015; Yuen et al. 2015) using Complete Genomics’ genome data have shown the DNMs can be detected with a low false positive error rate using appropriate filters (Methods and Materials and Supplementary Methods). Following similar analysis steps, we found approximately 65, 65, and 50 DNMs per fetal genome in the STD libraries from a cell pellet, in the cfDNA libraries, and in the LFR libraries, respectively (Table S3). Pairwise comparisons between cfDNA and cell pellet libraries demonstrated that approximately 88% of DNMs were shared between libraries for each amniocentesis sample (Fig. 2c).

In order to determine the accuracy of DNM calls and confirm that DNMs identified in the fetal genome were present in the child’s genome we contacted and collected buccal samples from 13 of the study participants. Potential DNMs were randomly selected for confirmation and several hundred base pairs surrounding each candidate DNM were amplified by PCR. 175 regions were successfully amplified and Sanger sequenced. Of these, 162 (92.5%) were found to harbor the potential DNM (Table S4). Candidates shared between two replicate libraries had a much higher confirmation rate (99.1%), in agreement with inherited variant rates (Fig. 2) and suggesting that using replicate libraries to confirm inherited or *de novo* variant calls is a robust method of evaluation. Potential DNMs were further confirmed to be true DNMs by Sanger sequencing the DNA of each parent. Only 3 of the 89 potential DNMs for which Sanger sequencing was successful for both parents were found to be inherited (Table S4). This suggests the overall false negative rate of our sequencing process is quite low (~3.4%).

The average age of mothers and fathers in our cohort at the time of the amniocentesis procedure was 37.8 and 43.3, respectively (Table S2). It has previously been shown that older fathers contribute a higher number of DNMs to their children than younger fathers (Kong et al. 2012; Jiang et al. 2013). To examine if this correlation could be seen in our cohort we plotted the total number of DNMs by maternal and paternal age. This resulted in the previously described pattern of an increase in the total number of DNMs with increasing paternal age. In our cohort paternal age contributed approximately 1.3 DNMs per additional year of age (Fig. 3a). Plotting the same data by maternal age did not show the same pattern; instead older mothers appeared to contribute less DNMs to their children (Fig. 3b). However, this may be due to the small sample size and the trend in our cohort that the oldest fathers tended to have the youngest wives (Fig. S1). Analysis of the base spectrum of DNMs did not differ significantly from that of inherited variants (Fig. S2). For those samples with LFR data the parental origin of most DNMs could be determined. This analysis showed the expected pattern of approximately 1.6X more DNMs from the father (Table S5).

**Figure 3.**
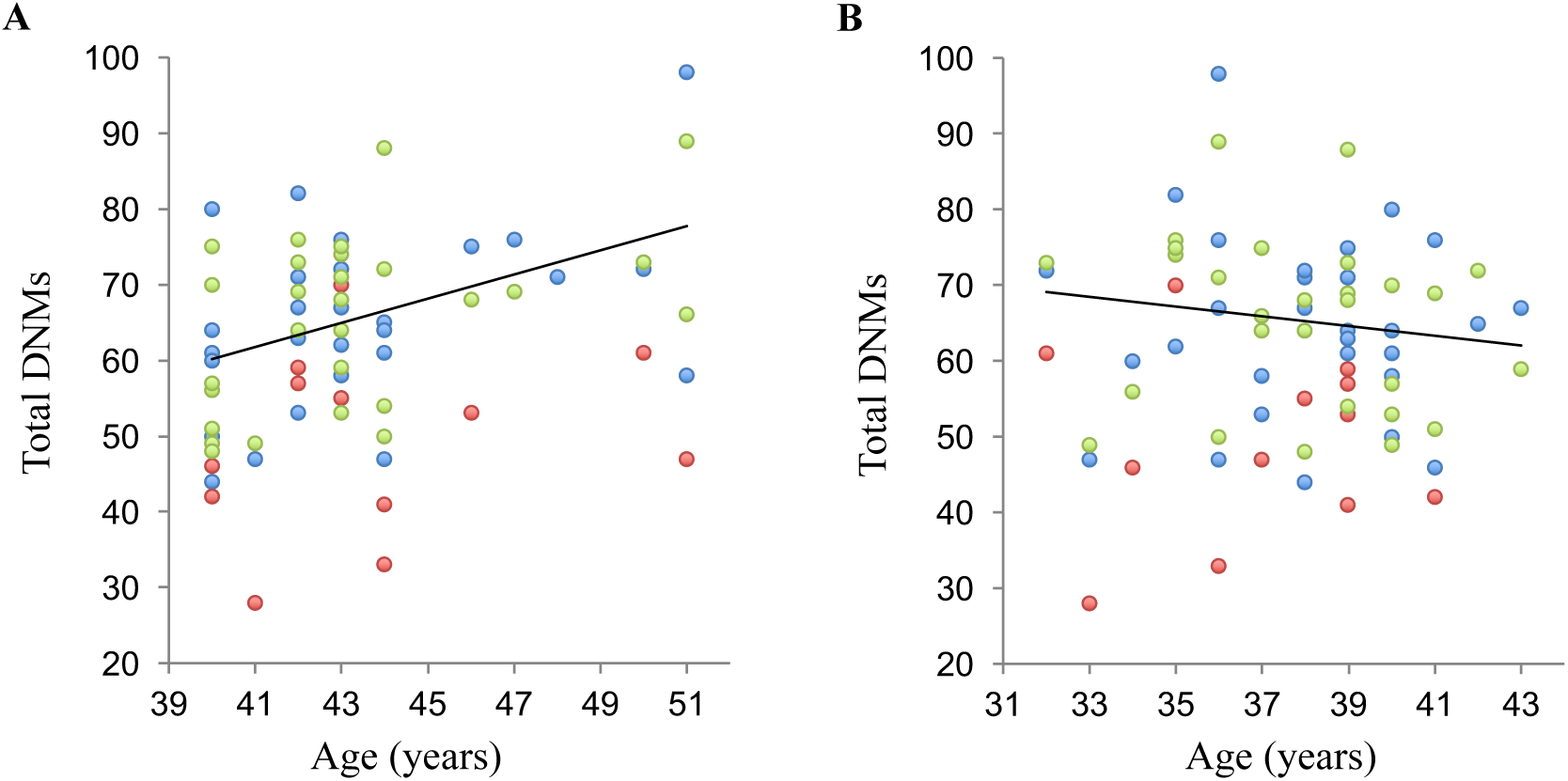
Parental age versus fetal DNMs. Paternal (**A**) and maternal (**B**) age were plotted on the x-axis versus total number of DNMs on the y axis. Each dot represents a fetal genome library. Blue diamonds denote libraries from cell pellets, red diamonds denote LFR libraries, and green diamonds represent libraries made from cfDNA. A trend line is plotted on each graph.

### Detection of copy number variants (CNVs) and structural variants (SVs)

Most of the women in this study were referred for an amniocentesis as a result of a positive noninvasive prenatal test (NIPT) for abnormal chromosomal copy number. For our WGS test to be useful, we should be able to detect these large-scale changes in structure and chromosomal copy number. To determine this we first compared our read coverage across all chromosomes to karyotyping results from the amniocentesis procedure. Two fetuses were found to carry an extra copy of chromosome 21 by karyotyping, this was also confirmed in our read coverage data for libraries made from cfDNA and cell pellet DNA (Fig. S3). In addition, there we three fetal genomes with know benign polymorphisms in heterochromatin and satellite DNA that were poorly covered by our WGS reads. This suggests that for many types of structural changes karyotyping will be necessary until WGS can be improved in these difficult to sequence parts of the genome.

However, many smaller changes (< 1 Mb) are difficult for karyotyping or array CGH to detect, but should be much easier for WGS. It is difficult to know what the ground truth is for CNVs and SVs in each genome, but having multiple replicates for each fetal sample and parental genome data enables reproducibility of our assay. As with small variants, we compared CNVs and SVs between replicate libraries and also compared them to parental genome calls (Tables S6 and S7). Over 94% of CNVs and 96% of SVs were found in at least one of the parents (Fig. 4a). In addition, ~66% of CNVs and ~74% of SVs overlapped with CNVs and SVs identified as part of the 1KG project (Sudmant et al. 2015) (Fig. 4b). Of those CNVs and SVs that were inherited, 97% and 85% were called between replicate libraries, respectively (Fig. 4c). In general, this demonstrates a high level of reproducibility and suggests that most CNV/SV calls are true positives.

**Figure 4.**
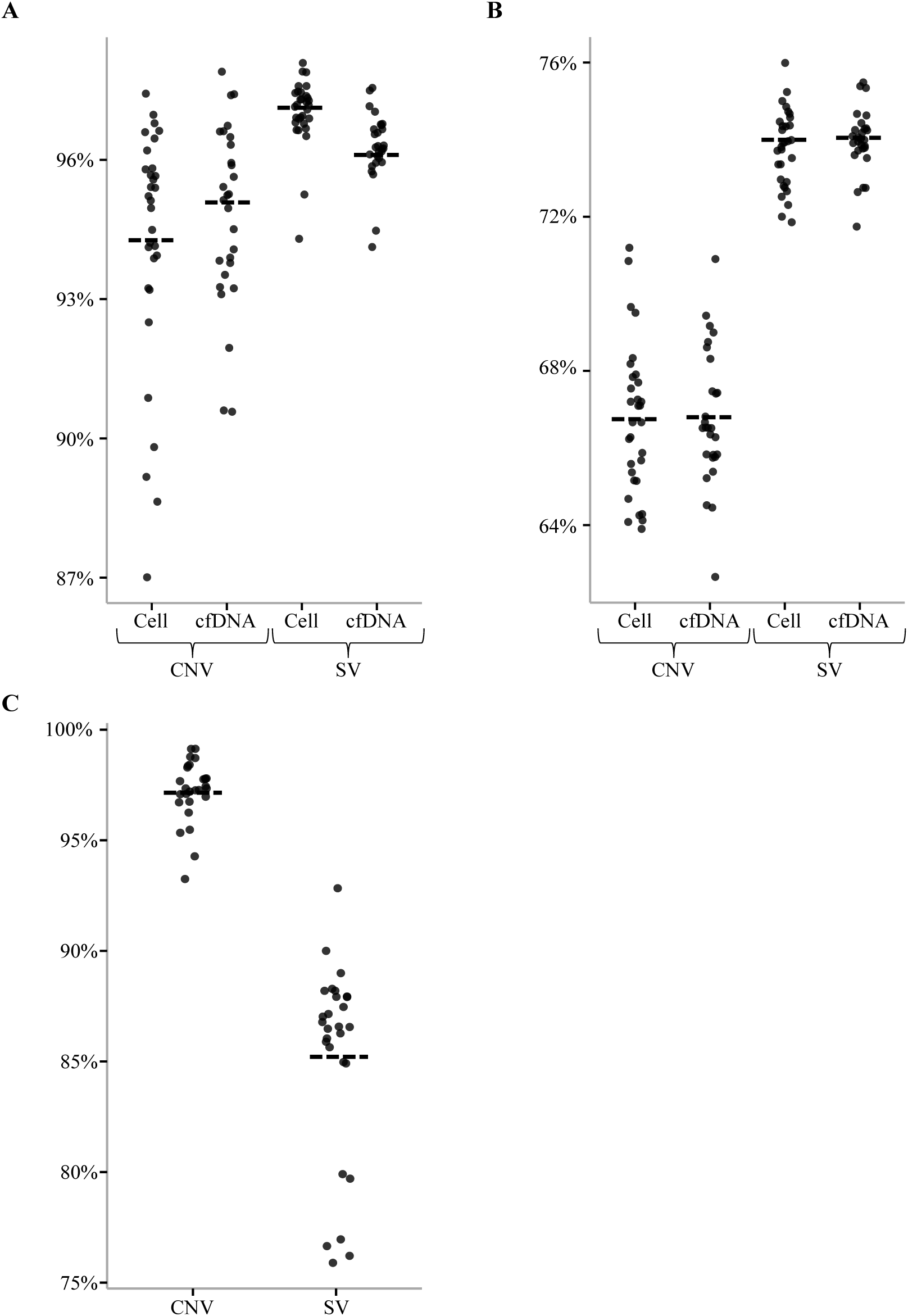
CNV/SV library concordance. The percent of CNVs and SVs called by STD libraries from the cell pellet (Cell) or amniotic fluid (cfDNA) and at least one parent (**A**) were plotted. The percent of CNVs and SVs called in fetal genomes and also called by the 1KG project was also plotted (**B**). Of those CNVs and SVs called by at least one parent the percent shared between the Cell and cfDNA libraries was plotted (**C**).

A total of 8 *de novo* CNV/SVs of greater than 1 kb were identified within the fetal genomes excluding trisomy 21 within the fetal genomes from families 21 and 22 (Tables S8 and S9). The largest identified was 14 kb. Based on previous studies, *de novo* CNVs larger than 100 kb are rare in healthy individuals (Sebat et al. 2007; Xu et al. 2008; Conrad et al. 2010; Itsara et al. 2010; Oskoui et al. 2015; Acuna-Hidalgo et al. 2016).

### Analysis of disease related genes

There is a growing list of databases that track the association of specific variants with disease. We searched our list of variants for entries in Clinvar (Landrum et al. 2014), a well know database for associating genomic variants with disease. On average, each fetal genome contained ~1 pathogenic or likely pathogenic variant with assertion criteria and no conflicting interpretations. Most of the potentially disease-causing variants appear to act in a recessive manner (Table S10) and no homozygous or compound heterozygous variants with these criteria were discovered. However, this means on average each child is a carrier for a potentially serious disease. We identified variants to such diseases as severe combined immunodeficiency, limb-girdle muscular dystrophy, and a predisposition to breast cancer. In addition, we found 6 children with different autosomal recessive deafness carrier alleles in the genes *GJB2* and *TMPRSS3*; these alleles are known to be more prevalent in Asian populations. None of the DNMs identified in our study were found in Clinvar.

To further analyze potential disease-causing variants that were not found in Clinvar we determined the Combined Annotation Dependent Depletion (CADD) (Kircher et al. 2014), SIFT (Kumar et al. 2009), and Polyphen2 (Adzhubei et al. 2010) scores for all rare coding inherited variants (Table S11) and DNMs (Table 1). We also used the ExAC (Lek et al. 2016) database to identify those genes with high pLI and missense Z-scores with variants and/or CNVs/SVs (Tables S8-S9). As a control these steps were repeated on the genomes of healthy Asian participants of the Personal Genome Project (PGP) (Mao et al. 2016). Based on this analysis the majority of variants appeared to be benign (Fig. 5). There were, however, a small number that merited further examination based on their scores, notably a detrimental DNM in *CHD8* in the fetal genome of FAM12. Mutations in this gene have recently been described as being one of the more common causes of ASD and define a particular subtype of the disease (Bernier et al. 2014). Contact of the family of this now 2-year old boy revealed that he does show at least one of the common phenotypes, macrocephaly, but at this time he does not show nor has he been evaluated for symptoms of ASD. In the fetal genome of FAM26 two different heterozygous missense variants, one from each parent, were identified in the gene *LRP1* (Table S11). Both are predicted to be detrimental by Polyphen2 and SIFT and have CADD scores above 24. Both variants are listed in ExAC, but are rare, one having been found in only two individuals in the database. In addition, the missense Z-score for this gene is 10.62 suggesting that it is highly intolerant to variation. Variants in this gene have been associated with Keratosis pilaris atrophicans, a skin disease that isn’t expected to severely affect the health of this child, however no additional information about whether this child shows any symptoms of the disease are available. The remainder of the children, as predicted by this genetic screen, have not been reported to have any serious illness.

**Figure 5.**
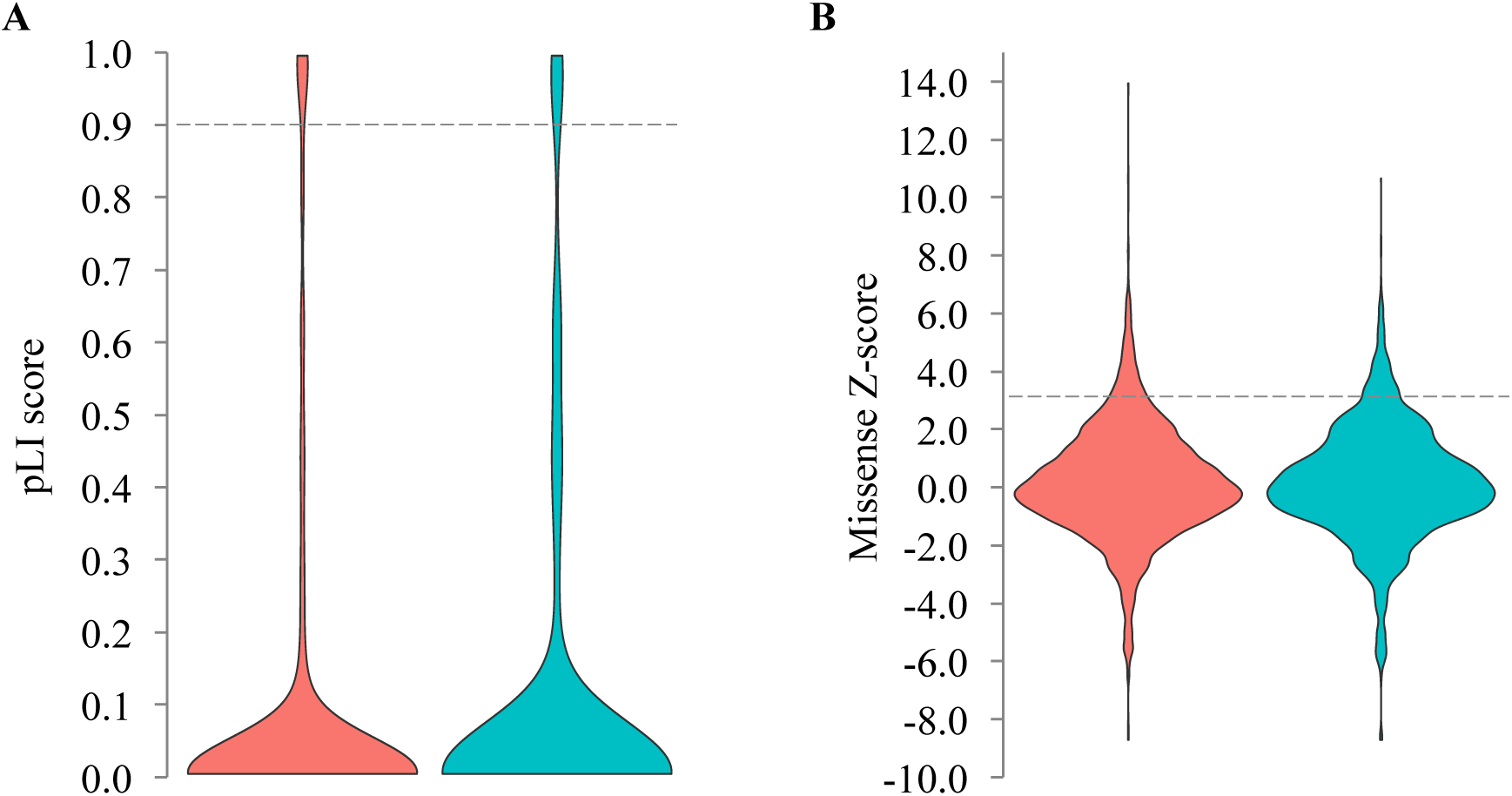
Gene pLI and missense Z-scores. The pLI scores (A) and missense Z-scores (B) from ExAC for genes with rare (MAF <= 0.01) protein truncating (frameshift, nonsense, splice site disrupting) and missense variants, respectively, were plotted. Amniocentesis samples are plotted in red and Asian genomes from the PGP project are plotted in green. The dotted line in each plot represents the level at which a gene is considered highly intolerant to variation.

**Table 1.**
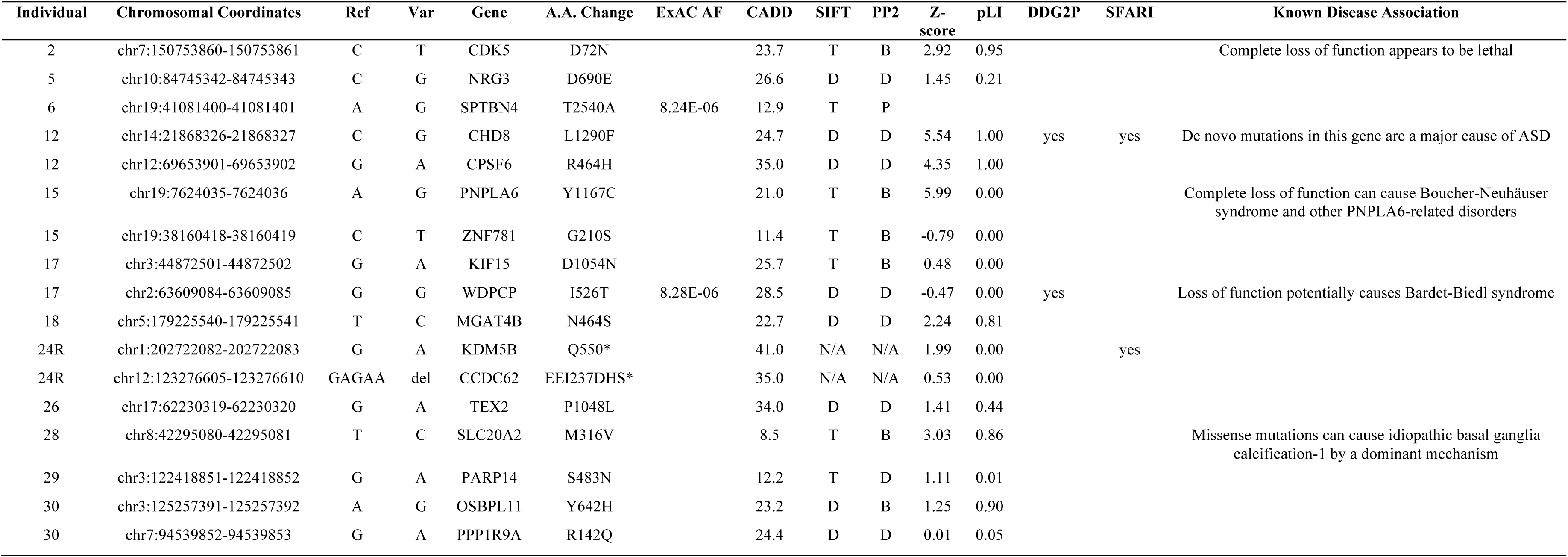
Coding DNMs. Coding DNMs in each fetal genome were annotated using publically available databases. For each DNM the fetal genome in which it was found (Individual), the genomic location it was found (Chromosomal Coordinates), the reference base at that position (Ref), the *de novo* base change at that position (Var), the gene it was found in (Gene), the amino acid change in the protein (A.A. Change), the frequency of this change in a database of over 100,000 exomes (ExAC Frequency), the CADD score (CADD), the likelihood that the change affects the protein function as determined by SIFT (SIFT) and Polyphen2 (PP2), the missense Z-score of the gene (Missense Score), the pLI score (pLI), if the gene was found to be associated with ID/DD (DDG2P) or autism (SFARI), and if the gene is known to be associated with a disease (Known Disease Association) are listed.

### Analysis of variants in genes targeted by drugs

Apart from diagnosing serious diseases there are other phenotypes that are important to identify in these fetal genomes. A tragic example is the case of a child who died from respiratory depression associated with excessive levels of morphine in the blood after elective adenotonsillectomy. It was later determined he had a duplication of *CYP2D6*, making him an ultrafast metabolizer of codeine; thus, the increased levels of its metabolite, morphine. He would have likely been prescribed a different dose or drug for pain management had this information been know (Ciszkowski et al. 2009). WGS analysis of amniotic material could potentially eliminate these types of severe interactions between a drug and an individual’s genetics from birth.

Each fetal genome in our study was analyzed against a list of potential drug interactions cataloged in the DrugBank database (Law et al. 2014). This resulted in over 400 coding variants per fetal genome in genes that are known targets of drugs. Analysis of Asian genomes and other ethnic groups from the 1KG project resulted in a similar number of coding drug target variants (Fig. S4). The vast majority of these variants would be unexpected to alter the protein product of these genes in such a way as to cause a serious adverse drug reaction. However, we discovered 381 instances of a variant with a low frequency in the population that resulted in complete loss of one copy of a gene listed in the DrugBank database in at least one of our fetal genomes. Again, it is unclear what affect, if any, these variants would have and improvements in our understanding of the interaction between drugs and specific variants will be necessary before this type of data can be fully utilized.

Currently, there are a few well-known gene-drug interactions that we can investigate. Specifically, the cytochrome 450 (CYP450) family involved with metabolizing most drugs and the genes involved in severe reactions to anesthesia. The results of this analysis are summarized in Table 2. Importantly, we discovered a large number of these children had at least one copy of an inactive or reduced activity CYP450. There are a number of drugs, such as warfarin, where dosing would be altered based on this information. In addition, we identified 4 rare damaging variants in *RYR1* and one in *CACNA1S.* Variants in these genes have been associated with malignant hyperthermia (MH), a serious and sometimes fatal response to anesthesia (Rosenberg et al. 2015). While it is unlikely that all 5 individuals are at risk for MH, this information would alert an anesthesiologist to utilize additional precautions and avoid MH triggering medications during the management of anesthetic care. A caffeine halothane contracture test on a muscle biopsy would also be recommended to confirm MH.

**Table 2.** Inherited variants in drug response genes. Well known gene drug interactions were examined. For each variant in a gene with certain alleles that are known to cause an interaction with a certain drug or class of drugs the fetal genome in which it was found (Individual), the genomic location it was found (Chromosomal Coordinates), the reference base at that position (Ref), the variant base at that position (Var), the dbSNP identifier (dbSNP), the gene it was found in (Gene), the amino acid change in the protein (A.A. Change), the frequency of this change in a database of over 100,000 exomes (ExAC Frequency), the allele frequency in a database of over 15,000 genomes (gnomAD AF), the CADD score (CADD), the likelihood that the change affects the protein function as determined by SIFT (SIFT) and Polyphen2 (PP2), and the known drug interaction (Drug Interaction) are listed.

|  | Individual | Chromosomal Coordinates | Ref | Var | dbSNP | Gene | A.A. Change | ExAC AF | gnomAD AF | CADD | SIFT | PP2 | Drug Interaction |
| --- | --- | --- | --- | --- | --- | --- | --- | --- | --- | --- | --- | --- | --- |
| Rare | 3 | chr19:38976581-38976582 | C | T | rs202225176 | RYR1 | P1763S | 6.60E-05 | 1.62E-04 | 22.9 | T | D | Some variants in this gene can cause malignant hyperthermia in response to some anesthesia |
|  | 6 | chr19:39010063-39010064 | C | T |  | RYR1 | P3410L | 2.47E-05 | 1.29E-04 | 25.8 | D | D |  |
|  | 10 | chr19:39008322-39008323 | G | A |  | RYR1 | R3337Q |  |  | 23.2 | D | D |  |
|  | 16 | chr19:38976507-38976508 | C | A |  | RYR1 | T1738K |  |  | 25.5 | T | D |  |
|  | 22 | chr1:201019609-201000000 | T | C |  | CACNA1S | N1383S | 8.24E-06 |  | 26.1 | D | D |  |
|  | 26 | chr22:42525034-42525035 | C | T | rs5030865 | CYP2D6 | G169R | 9.52E-04 | 2.93E-04 | 24.4 | D | D | This variant is part of CYP2D6*14 haplotype, it has no activity and can affect the dosage of many different drugs |
| Common | 3 hets | chr10:96741052-96741053 | A | C | rs1057910 | CYP2C9 | I359L | 0.064 | 0.048 | 20.4 | D | B | This variant is part of the CYP2C9*3 haplotype, it has low activity and can affect the dosage of many different drugs |
|  | 10 hets and 10 homs | chr22:42526693-42526694 | G | A | rs1065852 | CYP2D6 | P34S | 0.204 | 0.192 | 24.9 | D | D | This variant is part of the CYP2D6*10 haplotype, it has low activity and can affect the dosage of many different drugs |
|  | 13 hets and 3 homs | chr7:99270538-99270539 | C | T | rs776746 | CYP3A5 | Disrupt |  | 0.265 | 3.375 |  |  | This variant is part of the CYP3A5*3 haplotype, it has no activity and can affect the dosage of many different drugs |

## Discussion

In this study, we demonstrate for the first time the complete WGS analysis of amniotic samples from pregnant women. We show that up to 97% of the fetal genome can be confidently called using either DNA isolated from the fetal cell pellet or the amniotic fluid with virtually no difference in quality or coverage between the two sources. This is an important discovery as the leftover amniotic fluid is considered a waste product suggesting that WGS could be added to amniocentesis testing without interfering with the current standard of care tests. We also demonstrate that LFR libraries of high quality can be made from DNA isolated from the cell pellet allowing for haplotyping in these samples. We also discovered within the fetal genome almost all CNVs and SVs found in the parental genomes and many of these also overlapped with 1KG project samples. We identify 65 DNMs per genome from both cell pellet and cfDNA sources, in agreement with previous studies (Acuna-Hidalgo et al. 2016), and show that most of these are shared between the two libraries. In our data we find a previously described (Kong et al. 2012; Jiang et al. 2013) trend towards more DNMs in the genomes of those fetus’ with older fathers. Importantly, we find that over 92% of the DNMs identified by sequencing a single library from either the cell pellet or cfDNA exist in the newborn child, proving that this type of analysis is accurate and that the fetal genome is sufficiently predictive of the genome of the child.

In this cohort we discovered a single fetus with a DNM in *CHD8.* At 2 years of age it is already evident that this child has macrocephaly. Importantly, damaging DNMs in *CHD8* are one of the most common causes of simplex cases of ASD and 80% of individuals with ASD and a DNM in *CHD8* also display macrocephaly (Bernier et al. 2014). This suggests that the child in our study should be monitored for development of ASD. We also identified a child with compound heterozygous detrimental variants in *LRP1*, which could cause Keratosis pilaris atrophicans, although we were unable to obtain any additional information about the health of this child beyond that she was born healthy. We also found that many of the children in this study are carriers for a severe disease and importantly identified many that could have potential drug dosage issues due to reduced CYP450 activity. Finally, we identified 5 individuals we would consider to be at risk for MH due to rare variants in *RYR1* and *CACNA1S.* For those individuals, we would suggest that they undergo additional testing before being given general anesthesia or avoid anesthetic medications that are contraindicated for MH susceptible patients.

Importantly, through this analysis we show that much more information can be acquired from a routine amniocentesis procedure. At the current cost of about $1,000 we suggest the process we have described here should be considered as an additional analysis that can augment current karyotyping methods. This type of additional information has the potential to identify many of the causes of serious birth defects that currently go undetected.

## Materials and Methods

### Sample collection and processing

The institutional review boards of BGI-Shenzhen and the Peking University Shenzhen Hospital provided approval for this study. For each amniocentesis procedure 4 ml of amniotic fluid were sampled and frozen. Frozen amniotic fluid was thawed and centrifuged at 500X g for 5 minutes to pellet cells. The pellet was washed in PBS, followed by cell lysis, and purification by dialysis using a RecoverEase DNA isolation kit (Agilent Technologies, Santa Clara, CA). Approximately 100-1000 ng of high molecular weight genomic DNA were collected from each sample. Samples were concentrated using Microcon 30 kDa columns (Merck Millipore, Billerica, MA) to 105 ul. 5 ul and 100 ul were used for LFR library (Peters et al. 2012) and standard library construction (Drmanac et al. 2010), respectively, as previously described.

cfDNA was isolated from 3 ml of the remaining amniotic fluid supernatant from each sample using the QIAamp Circulating Nucleic Acid Kit (Qiagen, Hilden, Germany) and following the manufacturer’s protocol. Samples were eluted in 40ul TE buffer, which yielded between 80-1000 ng of DNA. Samples were brought to 100 ul with the addition of TE buffer and sheared by an E220 instrument (Covaris, Woburn, MA). Sheared samples were processed directly following a modified version of Complete Genomics’ (Mountain View, CA) standard library construction (Drmanac et al. 2010). Due to low starting material several purification steps were eliminated.

High molecular weight DNA was isolated from parental blood samples using a dialysis method as previously described (Peters et al. 2012) and processed using Complete Genomics’ standard library process (Drmanac et al. 2010). Both standard and LFR libraries were sequenced on Complete Genomics’s nanoarray platform. Sequence read mapping and variant calling were performed using Complete Genomics’s custom analysis pipeline(Carnevali et al. 2012). A full description of variant annotation and other analyses using whole genome data can be found in the supplementary materials.

### DNM analysis

DNMs in each fetal genome were first identified using the calldiff algorithm in CGA™ Tools (http://www.completegenomics.com/public-data/analysis-tools/cgatools/). Potential DNMs were screened against databases from the 1,000 genomes project (1KG) (Genomes Project et al. 2015), the Personal Genome Project (PGP) (Mao et al. 2016), the Wellderly Project (Erikson et al. 2016), and a Complete Genomics internal variant database to remove any variants that were false negatively called in the parental genomes.

### Sanger sequencing confirmation

To confirm DNMs buccal samples were collected from newborns using Omni swabs (GE Healthcare Life Sciences, Chicago, IL) and DNA was isolated using a QIAamp DNA Mini kit (Qiagen, Hilden, Germany) following the manufacturer’s protocol. High molecular weight DNA from the parents isolated for standard library construction and NA12878 were used to confirm DNMs were truly de novo and not false negatively called in the parents. PCR primers were designed to encompass approximately 250 base pairs on either side of the candidate DNM using Primer3 (Untergasser et al. 2012). Forward and reverse primers contained M13 forward and reverse primer sequences, respectively, to enable common primers to be used in Sanger sequencing. 1 ng of genomic DNA was used per PCR using AccuPrime *Taq* DNA polymerase (Thermo Fisher Scientific, Waltham, MA) following the manufacturer’s protocol. Sanger sequencing with both forward and reverse M13 primers was performed by McLab (South San Francisco, CA) and analyzed using Mutation Surveyor (SoftGenetics, State College, PA) with manual inspection of all results.

### Data access

Reads and mappings data have been submitted to the database of Genotypes and Phenotypes (dbGaP, http://www.ncbi.nlm.nih.gov/gap/) under accession ID phs001283.v1.p1.

## Acknowledgments

We would like to acknowledge the ongoing contributions and support of all Complete Genomics and BGI-Shenzhen employees, in particular the many highly skilled individuals that work in the libraries, reagents, and sequencing groups that make it possible to generate high quality whole genome data. This work was supported in part by the Shenzhen Municipal Government of China Peacock Plan NO.KQTD20150330171505310. Employees of BGI and Complete Genomics have stock holdings in BGI.

## Author contributions

B.A.P., R.D., and F.C. conceived the study. Y.D., W.X., and F.C. collected the amniocentesis samples. B.A.P., R.C., and R.Y.Z. developed the lab processes and made the libraries for sequence analysis. Q.M., N.G., Z.L., H.X., Q.S., E.E.P., and B.A.P performed analyses. B.A.P., W.X., F.C., and R.D. coordinated the study. B.A.P., R.C., and Q.M. wrote the paper. All authors contributed to revision and review of the manuscript.

## Supplemental Materials

Supplemental Methods

Supplemental Table S1. Summary statistics of all genomes.

Supplemental Table S2. Library IDs of each sample.

Supplemental Table S3. DNMs.

Supplemental Table S4. DNMs confirmed by Sanger sequencing.

Supplemental Table S5. LFR phasing of DNMs.

Supplemental Table S6. CNV counts

Supplemental Table S7. SV counts

Supplemental Table S8. CNVs

Supplemental Table S9. SVs

Supplemental Table S10. Clinvar variants

Supplemental Table S11. Rare variants

Supplemental Fig. S1. Paternal versus Maternal Age.

Supplemental Fig S2. Inherited and *de novo* single base spectrum changes.

Supplemental Fig S3. Chromosomal copy number analysis.

Supplemental Fig. S4. Gene drug interactions.

Supplemental Fig. S5. PLINK analysis.

Supplemental Fig. S6. Principal component analysis of samples.

## References

Acuna-Hidalgo R, Veltman JA, Hoischen A. 2016. New insights into the generation and role of de novo mutations in health and disease. Genome biology 17(1): 241.

Adzhubei IA, Schmidt S, Peshkin L, Ramensky VE, Gerasimova A, Bork P, Kondrashov AS, Sunyaev SR. 2010. A method and server for predicting damaging missense mutations. Nat Methods 7(4): 248-249.

Al Turki S, Manickaraj AK, Mercer CL, Gerety SS, Hitz MP, Lindsay S, D’Alessandro LC, Swaminathan GJ, Bentham J, Arndt AK et al. 2014. Rare variants in NR2F2 cause congenital heart defects in humans. Am J Hum Genet 94(4): 574-585.

Bernier R, Golzio C, Xiong B, Stessman HA, Coe BP, Penn O, Witherspoon K, Gerdts J, Baker C, Vulto-van Silfhout AT et al. 2014. Disruptive CHD8 mutations define a subtype of autism early in development. Cell 158(2): 263-276.

Carnevali P, Baccash J, Halpern AL, Nazarenko I, Nilsen GB, Pant KP, Ebert JC, Brownley A, Morenzoni M, Karpinchyk V et al. 2012. Computational techniques for human genome resequencing using mated gapped reads. J Comput Biol 19(3): 279-292.

Ciszkowski C, Madadi P, Phillips MS, Lauwers AE, Koren G. 2009. Codeine, ultrarapid-metabolism genotype, and postoperative death. N Engl J Med 361(8): 827-828.

Conrad DF, Pinto D, Redon R, Feuk L, Gokcumen O, Zhang Y, Aerts J, Andrews TD, Barnes C, Campbell P et al. 2010. Origins and functional impact of copy number variation in the human genome. Nature 464(7289): 704-712.

de Ligt J, Willemsen MH, van Bon BW, Kleefstra T, Yntema HG, Kroes T, Vulto-van Silfhout AT, Koolen DA, de Vries P, Gilissen C et al. 2012. Diagnostic exome sequencing in persons with severe intellectual disability. N Engl J Med 367(20): 1921-1929.

De Rubeis S, He X, Goldberg AP, Poultney CS, Samocha K, Ercument Cicek A, Kou Y, Liu L, Fromer M, Walker S et al. 2014. Synaptic, transcriptional and chromatin genes disrupted in autism. Nature.

Deciphering Developmental Disorders S. 2015. Large-scale discovery of novel genetic causes of developmental disorders. Nature 519(7542): 223-228.

Drmanac R, Sparks AB, Callow MJ, Halpern AL, Burns NL, Kermani BG, Carnevali P, Nazarenko I, Nilsen GB, Yeung G et al. 2010. Human genome sequencing using unchained base reads on self-assembling DNA nanoarrays. Science 327(5961): 78-81.

Epi KC, Epilepsy Phenome/Genome P, Allen AS, Berkovic SF, Cossette P, Delanty N, Dlugos D, Eichler EE, Epstein MP, Glauser T et al. 2013. De novo mutations in epileptic encephalopathies. Nature 501(7466): 217-221.

Erikson GA, Bodian DL, Rueda M, Molparia B, Scott ER, Scott-Van Zeeland AA, Topol SE, Wineinger NE, Niederhuber JE, Topol EJ et al. 2016. Whole-Genome Sequencing of a Healthy Aging Cohort. Cell.

Fromer M, Pocklington AJ, Kavanagh DH, Williams HJ, Dwyer S, Gormley P, Georgieva L, Rees E, Palta P, Ruderfer DM et al. 2014. De novo mutations in schizophrenia implicate synaptic networks. Nature 506(7487): 179-184.

Genomes Project C, Auton A, Brooks LD, Durbin RM, Garrison EP, Kang HM, Korbel JO, Marchini JL, McCarthy S, McVean GA et al. 2015. A global reference for human genetic variation. Nature 526(7571): 68-74.

Gilissen C, Hehir-Kwa JY, Thung DT, van de Vorst M, van Bon BW, Willemsen MH, Kwint M, Janssen IM, Hoischen A, Schenck A et al. 2014. Genome sequencing identifies major causes of severe intellectual disability. Nature 511(7509): 344-347.

Iossifov I, O’Roak BJ, Sanders SJ, Ronemus M, Krumm N, Levy D, Stessman HA, Witherspoon KT, Vives L, Patterson KE et al. 2014. The contribution of de novo coding mutations to autism spectrum disorder. Nature 515(7526): 216-221.

Itsara A, Wu H, Smith JD, Nickerson DA, Romieu I, London SJ, Eichler EE. 2010. De novo rates and selection of large copy number variation. Genome Res 20(11): 1469-1481.

Jiang YH, Yuen RK, Jin X, Wang M, Chen N, Wu X, Ju J, Mei J, Shi Y, He M et al. 2013. Detection of clinically relevant genetic variants in autism spectrum disorder by whole-genome sequencing. Am J Hum Genet 93(2): 249-263.

Kircher M, Witten DM, Jain P, O’Roak BJ, Cooper GM, Shendure J. 2014. A general framework for estimating the relative pathogenicity of human genetic variants. Nat Genet 46(3): 310-315.

Kong A, Frigge ML, Masson G, Besenbacher S, Sulem P, Magnusson G, Gudjonsson SA, Sigurdsson A, Jonasdottir A, Wong WS et al. 2012. Rate of de novo mutations and the importance of father’s age to disease risk. Nature 488(7412): 471-475.

Kumar P, Henikoff S, Ng PC. 2009. Predicting the effects of coding non-synonymous variants on protein function using the SIFT algorithm. Nat Protoc 4(7): 1073-1081.

Landrum MJ, Lee JM, Riley GR, Jang W, Rubinstein WS, Church DM, Maglott DR. 2014. ClinVar: public archive of relationships among sequence variation and human phenotype. Nucleic Acids Res 42(Database issue): D980-985.

Law V, Knox C, Djoumbou Y, Jewison T, Guo AC, Liu Y, Maciejewski A, Arndt D, Wilson M, Neveu V et al. 2014. DrugBank 4.0: shedding new light on drug metabolism. Nucleic Acids Res 42(Database issue): D1091-1097.

Lek M, Karczewski KJ, Minikel EV, Samocha KE, Banks E, Fennell T, O’Donnell-Luria AH, Ware JS, Hill AJ, Cummings BB et al. 2016. Analysis of protein-coding genetic variation in 60,706 humans. Nature 536(7616): 285-291.

Mao Q, Ciotlos S, Zhang RY, Ball MP, Chin R, Carnevali P, Barua N, Nguyen S, Agarwal MR, Clegg T et al. 2016. The whole genome sequences and experimentally phased haplotypes of over 100 personal genomes. Gigascience 5(1): 42.

McCarthy SE, Gillis J, Kramer M, Lihm J, Yoon S, Berstein Y, Mistry M, Pavlidis P, Solomon R, Ghiban E et al. 2014. De novo mutations in schizophrenia implicate chromatin remodeling and support a genetic overlap with autism and intellectual disability. Mol Psychiatry.

Michaelson JJ, Shi Y, Gujral M, Zheng H, Malhotra D, Jin X, Jian M, Liu G, Greer D, Bhandari A et al. 2012. Whole-genome sequencing in autism identifies hot spots for de novo germline mutation. Cell 151(7): 1431-1442.

O’Roak BJ, Vives L, Girirajan S, Karakoc E, Krumm N, Coe BP, Levy R, Ko A, Lee C, Smith JD et al. 2012. Sporadic autism exomes reveal a highly interconnected protein network of de novo mutations. Nature 485(7397): 246-250.

Oskoui M, Gazzellone MJ, Thiruvahindrapuram B, Zarrei M, Andersen J, Wei J, Wang Z, Wintle RF, Marshall CR, Cohn RD et al. 2015. Clinically relevant copy number variations detected in cerebral palsy. Nat Commun 6: 7949.

Peters BA, Kermani BG, Alferov O, Agarwal MR, McElwain MA, Gulbahce N, Hayden DM, Tang YT, Zhang RY, Tearle R et al. 2015. Detection and phasing of single base de novo mutations in biopsies from human in vitro fertilized embryos by advanced whole-genome sequencing. Genome Res 25(3): 426-434.

Peters BA, Kermani BG, Sparks AB, Alferov O, Hong P, Alexeev A, Jiang Y, Dahl F, Tang YT, Haas J et al. 2012. Accurate whole-genome sequencing and haplotyping from 10 to 20 human cells. Nature 487(7406): 190-195.

Purcell SM, Moran JL, Fromer M, Ruderfer D, Solovieff N, Roussos P, O’Dushlaine C, Chambert K, Bergen SE, Kahler A et al. 2014. A polygenic burden of rare disruptive mutations in schizophrenia. Nature 506(7487): 185-190.

Rosenberg H, Pollock N, Schiemann A, Bulger T, Stowell K. 2015. Malignant hyperthermia: a review. Orphanet J Rare Dis 10: 93.

Sanders SJ, Murtha MT, Gupta AR, Murdoch JD, Raubeson MJ, Willsey AJ, Ercan-Sencicek AG, DiLullo NM, Parikshak NN, Stein JL et al. 2012. De novo mutations revealed by whole-exome sequencing are strongly associated with autism. Nature 485(7397): 237-241.

Sebat J, Lakshmi B, Malhotra D, Troge J, Lese-Martin C, Walsh T, Yamrom B, Yoon S, Krasnitz A, Kendall J et al. 2007. Strong association of de novo copy number mutations with autism. Science 316(5823): 445-449.

Sudmant PH, Rausch T, Gardner EJ, Handsaker RE, Abyzov A, Huddleston J, Zhang Y, Ye K, Jun G, Hsi-Yang Fritz M et al. 2015. An integrated map of structural variation in 2,504 human genomes. Nature 526(7571): 75-81.

Untergasser A, Cutcutache I, Koressaar T, Ye J, Faircloth BC, Remm M, Rozen SG. 2012. Primer3--new capabilities and interfaces. Nucleic Acids Res 40(15): e115.

Veltman JA, Brunner HG. 2012. De novo mutations in human genetic disease. Nat Rev Genet 13(8): 565-575.

Xu B, Roos JL, Levy S, van Rensburg EJ, Gogos JA, Karayiorgou M. 2008. Strong association of de novo copy number mutations with sporadic schizophrenia. Nat Genet 40(7): 880-885.

Yang Y, Muzny DM, Reid JG, Bainbridge MN, Willis A, Ward PA, Braxton A, Beuten J, Xia F, Niu Z et al. 2013. Clinical Whole-Exome Sequencing for the Diagnosis of Mendelian Disorders. New England Journal of Medicine 369(16): 1502-1511.

Yuen RK, Thiruvahindrapuram B, Merico D, Walker S, Tammimies K, Hoang N, Chrysler C, Nalpathamkalam T, Pellecchia G, Liu Y et al. 2015. Whole-genome sequencing of quartet families with autism spectrum disorder. Nat Med 21(2): 185-191.

